## Supplementary File for "Phase transition of WTAP regulates m^6^A modification of interferon-stimulated genes"

**Extended Methods**

**Expression, purification, and interaction of mCherry-WTAP WT, 5ST-A/ST-D mutants, CFP-METTL3 and GFP-STAT1.**

Procedures of the expression of proteins in *E. coli* were referred to Hao Jiang et al (57). Recombinant mCherry-WTAP, CFP-METTL3 and GFP-STAT1 were cloned into pET-28A plasmid, while ST-D and ST-A mutants of WTAP were constructed by mutating the pET-28A-mCherry-WTAP plasmid using Muta-direct^TM^ Kit (Sbsgene, cat. #SDM-15). After transformed into *E.* *coli*, bacteria were cultured at 37℃ overnight until OD600 of 0.4 and 1 mM IPTG was used to induced the expression of mCherry-WTAP for 6 hours. After IPTG induction, bacteria were lysed by sonication with lysis buffer (50 mM NaH_2_PO_4_, 300 mM NaCl, pH 7.8). Lysates were centrifugated at 12000 g, 37°C for 10 minutes and supernatants were incubated with Ni-NTA beads (Thermo Fisher Scientific, cat. #R90101) overnight. Beads were then washed by washing buffer (50 mM NaH_2_PO_4_, 300 mM NaCl, 20 mM imidazole pH 7.8) and eluted by elution buffer (50 mM NaH_2_PO_4_, 300 mM NaCl, 500 mM imidazole pH 7.8). The protein solution was concentrated using Amicon® Ultra 30K device (Millipore, cat. #C134281) and concentration of protein was detected using Pierce™ BCA Protein Assay Kit (cat. #23225, Thermo Fisher Scientific) as manufacturer’s instructions.

For detection of interaction among mCherry-WTAP or its indicated mutants, CFP-METTL3 and GFP-STAT, purified proteins were mixed using physiological buffer (20 mM Tris-HCl, pH 7.5, 150 mM KCl, 10% PEG8000) in the final concentration of 10 μM and placed on the slides. Mixtures were incubated and imaged at 37^o^C in a live-cell-imaging chamber of Leica TCS-SP8 STED 3X confocal fluorescence microscope while images were analyzed by LAS AF software.

**Mass spectrometry.**

HEK 293T cells were transfected with plasmids expressing mCherry-WTAP. 24 hours after transfection, whole cell lysate was obtained using low-salt lysis buffer and immunoprecipitated using protein A/G beads together with anti-WTAP antibody at 4°C overnight. Beads were washed five times with low-salt lysis buffer and boiled at 100 °C for 5 minutes with 2×SDS loading buffer, followed with immunoblot assays described above. Protein bands were visualized by Coomassie Blue R-250 staining. Bands of around 44 kDa was obtained for analyzing phosphorylation sites of WTAP. Protein digestion was performed by trypsin. The digested peptides of each sample were desalted on C18 Cartridges (Empore™ SPE Cartridges C18 (standard density), bed I.D. 7 mm, volume 3 ml, Sigma), concentrated by vacuum centrifugation and reconstituted in 40 µl of 0.1% (v/v) formic acid.

LC-MS/MS analysis was performed by Applied Protein Technology (Shanghai, China) on a Q Exactive mass spectrometer (Thermo Scientific) that was coupled to Easy nLC (Proxeon Biosystems, now Thermo Fisher Scientific) for 120 min. The peptides were loaded onto a reverse phase trap column (Thermo Scientific Acclaim PepMap100, 100 μm*2 cm, nanoViper C18) connected to the C18-reversed phase analytical column (Thermo Scientific Easy Column, 10 cm long, 75 μm inner diameter, 3μm resin) in buffer A (0.1% Formic acid) and separated with a linear gradient of buffer B (84% acetonitrile and 0.1% Formic acid) at a flow rate of 300 nl/min controlled by IntelliFlow technology. The mass spectrometer was operated in positive ion mode. MS data was acquired using a data-dependent top10 method dynamically choosing the most abundant precursor ions from the survey scan (300–1800 m/z) for HCD fragmentation. Automatic gain control (AGC) target was set to 3e6, and maximum inject time to 10 ms. Dynamic exclusion duration was 40.0 s. Survey scans were acquired at a resolution of 70,000 at m/z 200 and resolution for HCD spectra was set to 17,500 at m/z 200, and isolation width was 2 m/z. Normalized collision energy was 30 eV and the underfill ratio, which specifies the minimum percentage of the target value likely to be reached at maximum fill time, was defined as 0.1%. The instrument was run with peptide recognition mode enabled. The MS raw data for each sample were combined and searched using the MaxQuant software for identification and quantitation analysis.

**Chromatin-immunoprecipitation (ChIP)-qPCR assay.**

The procedures were referred to the Rockland Chromatin Immunoprecipitation assay protocol. Protein A or G Sepharose were blocked by 5% BSA and 1 μg/mL HT-DNA for 2 hours, then 4 μg antibody was added to incubate for 2 hours. Relevant cells with indicated treatment were mixed with 1% formaldehyde and incubated at room temperature for 10 minutes, then mixed with 10% glycine from 1.375 M stock. The cells were washed with ice-cold PBS for 3 times, scaped and collected cells in PBS. After centrifuging at 1,000 rpm for 5 minutes in cold centrifuge, used 10 volume swelling buffer (25 mM Hepes, pH 7.8, 1.5 mM MgCl_2_ 10 mM KCl, 0.1% NP-40, 1 mM DTT, 0.5 mM PMSF, protease inhibitor cocktail) to resuspend pellet and incubated in ice for 10 minutes. The cells were ground 10-20 times, centrifuged at 2,000 rpm for 5 minutes. The pullets were resuspended in sonication buffer (50 mM Hepes, pH 7.9, 140 mM NaCl, 1 mM EDTA, 1% Triton X-100, 0.1% Na-deoxycholate, 0.1% SDS, 0.5 mM PMSF, protease inhibitor cocktail) to performed 9 times sonication for 10-20 seconds at 80% setting in sonicator. After centrifuging at 14,000 rpm for 15 minutes, the supernatants were collected, 10% input sample was divided, and the rest was co-incubated with indicated antibodies and beads by constant rotation in the cold room overnight.

The mixtures were centrifuged 6,000 rpm for 3 minutes and the beads were washed 2 times with 1 mL sonication buffer, 2 times with 1 mL wash buffer A (50 mM Hepes, pH 7.9, 500 mM NaCl, 1 mM EDTA, 1% Triton X-100, 0.1% Na-deoxycholate, 0.1% SDS, 0.5 mM PMSF, protease inhibitor cocktail), 2 times with 1mL wash buffer B (20 mM Tris, pH 8.0, 1 mM EDTA, 250 mM LiCl, 0.5% NP-40, 0.5% Na-deoxycholate, 0.5 mM PMSF, protease inhibitor cocktail), and 2 times with 1 mL TE buffer. 200 μL Elution buffer (50 mM Tris, pH 8.0, 1 mM EDTA, 1% SDS, 50 mM NaHCO_3_) were added to beads and incubated at 65℃ for 10 minutes. Centrifuged to collect the supernatant and elute the beads again, combined the eluates. 21 μL NaCl from 4 M stock were added to input (added elution buffer to 400 μL), IgG and IP samples which incubated at 65℃ overnight. 1 μL RNase A from 10 mg/mL were added and incubated at 37℃ for 1 hour, along with 4 μL EDTA from 0.5 M stock and 2 μL proteinase K from 10 mg/mL were added and incubated at 42℃ for 2 hours.

DNA was extracted once with phenol/chloroform/isoamylalcohol followed with centrifuging at 12,000 rpm for 2 minutes, the clean aqueous phase to new tube and once with chloroform/isoamylacohol followed with centrifuging at 12,000 rpm for 2 minutes. The samples were then added 1 μL glycogen from 20 mg/mL stock as well as 1 mL pre-cold EtOH, and then left to precipitate in -20℃ overnight. After centrifuging for 15 minutes, pullets were washed 1 time with 80% EtOH and resuspended with 20 μL TE buffer, followed with qPCR assay.

**RNA-immunoprecipitation (RIP) followed with qPCR assay.**

Approximately 1×10^7^ cells were differentiated and stimulated with 10 ng/mL IFN-β for 2 hours. After stimulation, cells were collected and lysed by polysome lysis buffer (100 mM KCl, 5 mM MgCl_2_, 10 mM HEPES (pH 7.0), 0.5% NP-40, 1 mM DTT, 100 units/mL recombinant RNase inhibitor (cat. #2313B, Takara), 400 μM vanadyl ribonucleoside complexes (VRC, cat. #S1402S, New England Biolabs), protease inhibitor cocktail (cat. #4906837001, Roche)). 50 μL protein A/G beads were pre-swelled by 5 times volume of 5% BSA and incubated with 4 μg indicated antibody overnight. After antibody incubation, beads were washed by NT2 buffer (50 mM Tris-HCl (pH 7.4), 150 mM NaCl, 1 mM MgCl_2_, 0.05% NP-40) for 5 times and incubated with cell lysate for 4 hours, while 5% of cell lysate was kept as input. After immunoprecipitation, precipitates were washed by NT2 buffer for 5 times and supplemented with 30 μg proteinase K 55°C for 30 minutes to release RNA-protein complex. Finally, RNA was isolated by TRIzol reagent and analyzed by qPCR assay as above.

**RNA sequencing (RNA-Seq) assay.**

For RNA-Seq assay, total RNA was isolated from cells using TRIzol reagent, and sequencing was performed by Sangon Biotech (Shanghai, China). Sample quality was assessed using a Bioanalyzer (Agilent 2100 Bioanalyzer). RNA-seq libraries of polyadenylated RNA were prepared using mRNA-seq V2 Library Prep Kit and sequenced on MGISEQ-2000 platform. All clean data were mapped to the human genome GRCh38 using STAR v2.7.9a with default parameters. Bam files were sorted by Samtools 1.9. Reads counts were summarized using the htseq-count tool as part of the HTSeq framework release v0.13.5 (https://htseq.readthedocs.io/). To identify DEGs between groups, DESeq2 was used to normalize reads counts and P-value < 0.05 and absolute logged fold-change ≥ 1 were determined as DEGs using Bioconductor clusterProfiler package v3.14.3 for the functional enrichment of DEGs. RNA-seq sequence density profiles were normalized using bamCoverage and visualized in IGV genome browser.

**MeRIP-Seq assay**.

For MeRIP-Seq, approximately 5×10^7^ cells were seeded and stimulated by 10 ng/mL IFN-β for 4 hours. After stimulation, total RNA was extracted using TRIzol reagent. The total RNA quality and quantity were analysis of Bioanalyzer 2100 and RNA 6000 Nano LabChip Kit (Agilent, CA, USA) with RIN number >7.0. Approximately more than 50 μg of total RNA was subjected to isolate poly (A) mRNA with poly-T oligo attached magnetic beads (cat. #61006, Invitrogen). Following purification, the poly(A) mRNA fractions is fragmented into ~100-nt-long oligonucleotides using divalent cations under elevated temperature. Then the cleaved RNA fragments were subjected to incubation for 2h at 4°C with m^6^A-specific antibody (cat.#202003, Synaptic Systems) in IP buffer (50 mM Tris-HCl, 750 mM NaCl and 0.5% Igepal CA-630) supplemented with BSA (0.5 μg/μL). The mixture was then incubated with protein-A beads and eluted with elution buffer (1×IP buffer and 6.7mM m^6^A). Eluted RNA was precipitated by 75% ethanol. Eluted m^6^A-containing fragments (IP) and untreated input control fragments are converted to final cDNA library in accordance with a strand-specific library preparation by dUTP method. The average insert size for the paired-end libraries was ~100±50 bp. And then we performed the paired-end 2×150 bp sequencing on an Illumina Novaseq 6000 platform at the LC-BIO Bio-tech ltd (Hangzhou, China) following the vendor's recommended protocol. For MeRIP-Seq data analysis, filtered reads were aligned to the human genome (hg38) using STAR with default settings and all non-unique mapped reads or PCR duplicates were removed. Next, we used exomePeak, an exome-based peak-calling R package(55), to detect significantly enriched m^6^A modification site (FDR < 0.05, fold enrichment $\geq$ 1.5) with default parameters, in which the changes in the expression level of m^6^A-modified gene in the control and treatment groups were considered. m^6^A sites were predicted using Deep-m^6^A, a software program for m^6^A site prediction(56). The consensus motif was determined using WebLogo v3.0. Peak browsing and representative snapshots capturing were performed using the Integrative Genomics Viewer (IGV2.4.10, Broad Institute)

**RNA decay assay.**

THP-1-derived macrophages were seeded with the density of 1×10^6^ cells/mL and pretreated with 10 ng/mL IFN-β for 4 hours for the induction of ISGs. For 1,6-hex treatment experiments, cells were pretreated with 10 ng/mL IFN-β, 5% 1,6-hex and 20 μg/mL digitonin for 2 hours and medium was replaced by fresh 1640 medium for 2 hours. After pretreatment, fresh medium containing 5 μM Actinomycin D was replaced and cells were collected at indicated time points followed with RNA extraction and qPCR assay as described above.

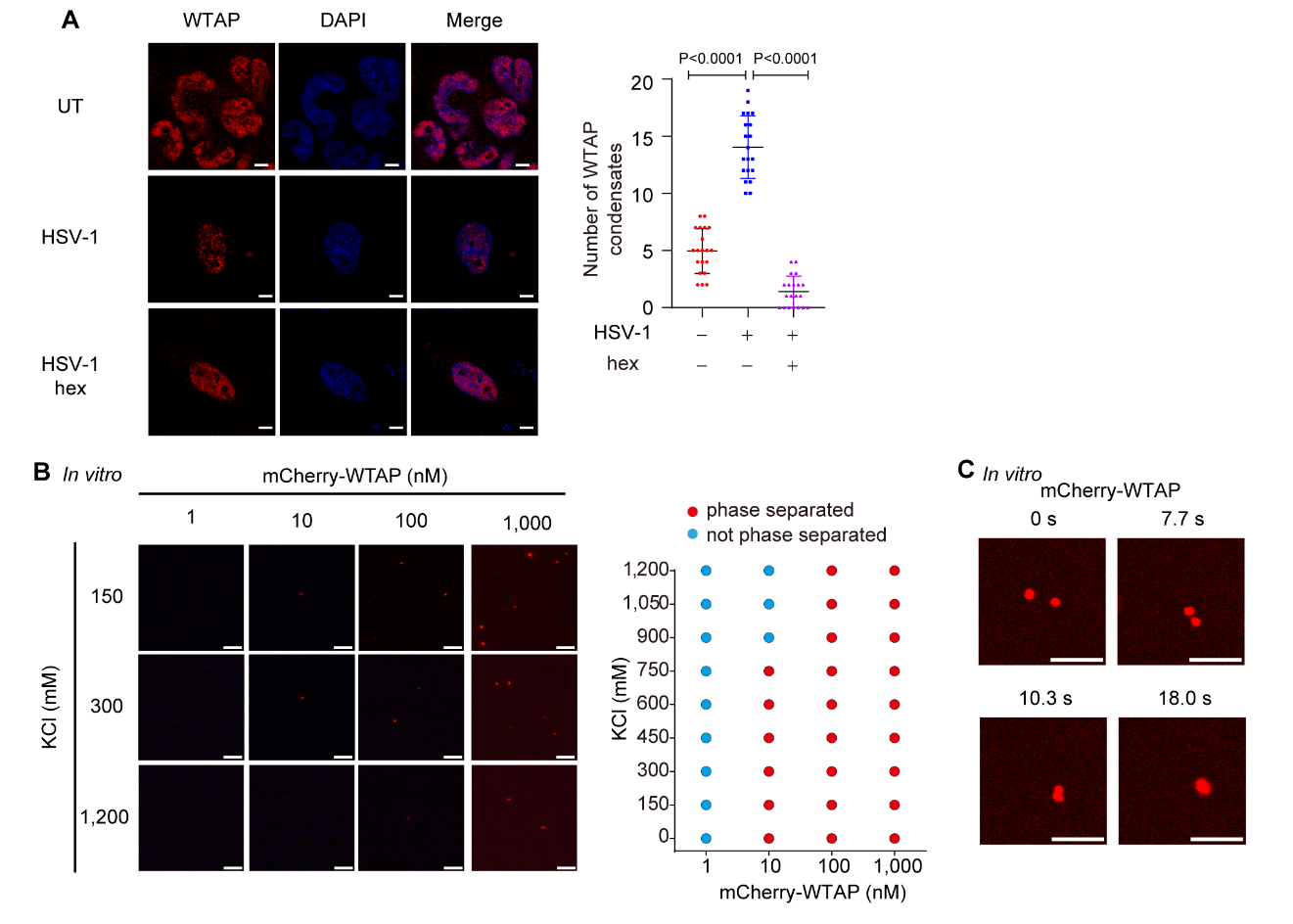

**Figure 1-figure supplement 1. WTAP undergoes phase separation*.***

(A) THP-1-derived macrophages were infected with Herpes Simplex Virus-1 (HSV-1) (m.o.i.=1) for 24 hours together with or without 5% 1,6-hexanediol (hex) and 10 μg/mL digitonin for 2 hours or left untreated (UT). WTAP were stained and imaged using confocal microscope. The number of WTAP condensates that diameter over 0.4 μm of n = 20 cells were counted through ImageJ and shown. Scale bars indicated 5 μm.

(B) Phase separation of recombinant mCherry-WTAP in different concentrations incubated with different concentrations of KCl were observed through confocal microscopy. Representative images were shown (left), and phase separation diagram was listed (right). Scale bars indicated 10 μm.

(C) Recombinant mCherry-WTAP (10 μM) was mixed with physiological buffer and incubated at 37 °C. Time-lapse microscopy of merging condensates was performed and representative images were shown. The beginning of our observation by confocal microscopy was defined as 0 second. Scale bars indicated 5 μm.

All error bars, mean values ± SD, P-values were determined by unpaired two-tailed Student’s *t*-test of n = 20 cells in (A). For (B-C), similar results were obtained for three independent biological experiments.

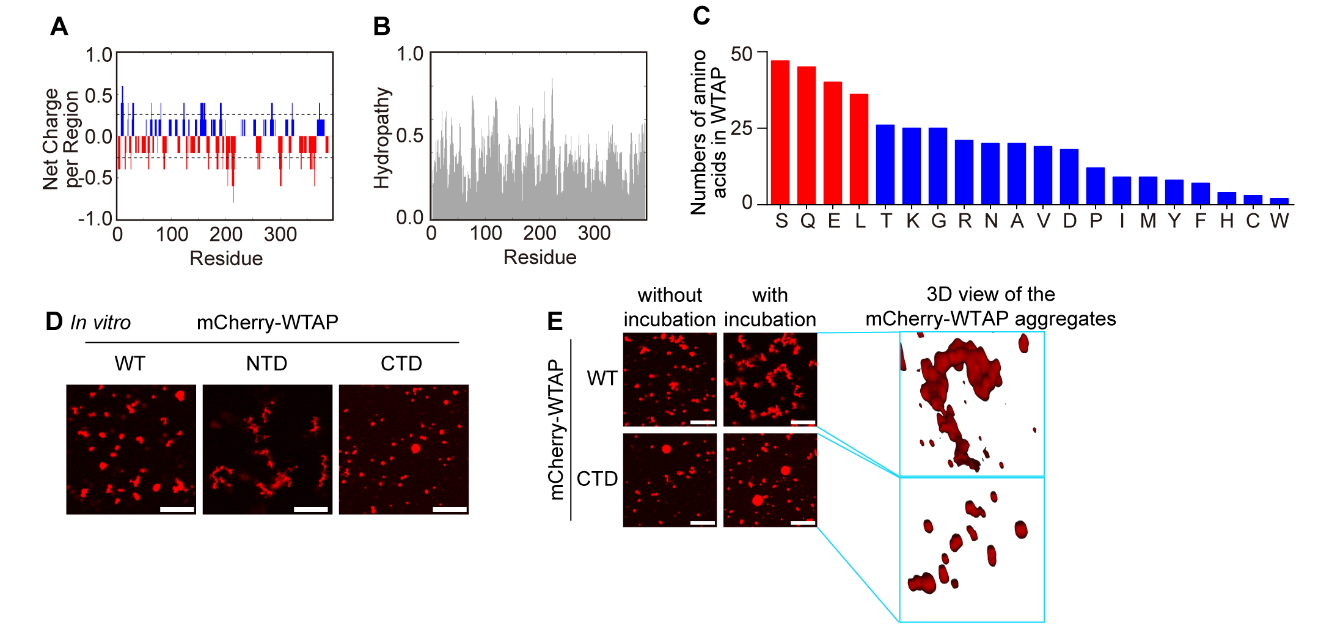

**Figure 1-figure supplement 2. Serine-rich CTD and glutamine-rich NTD of WTAP provide the potential of aggregates or liquid droplets of WTAP respectively.**

(A-B) Net charge per residue (A) and hydropathy (B) analysis of WTAP was performed through CIDER predictor (http://pappulab.wustl.edu/CIDER/analysis/).

(C) Numbers of different amino acids within WTAP was calculated and shown. Four abundant amino acids (Serine (S), glutamine (G), glutamic acid (E) and leucine (L)) were marked as red while the rest were labelled as blue.

(D) Phase separation of recombinant mCherry-WTAP WT, NTD and CTD (40 μM) were observed through confocal microscopy. Scale bars indicated 10 μm.

(E) Recombinant mCherry-WTAP (10 μM) was mixed with physiological buffer and incubated at 37 °C. Time-lapse microscopy of merging condensates were performed and representative images were shown. The beginning of our observation by confocal microscopy was defined as 0 second. Scale bars indicated 10 μm.

For (D-E), similar results were obtained for three independent biological experiments.

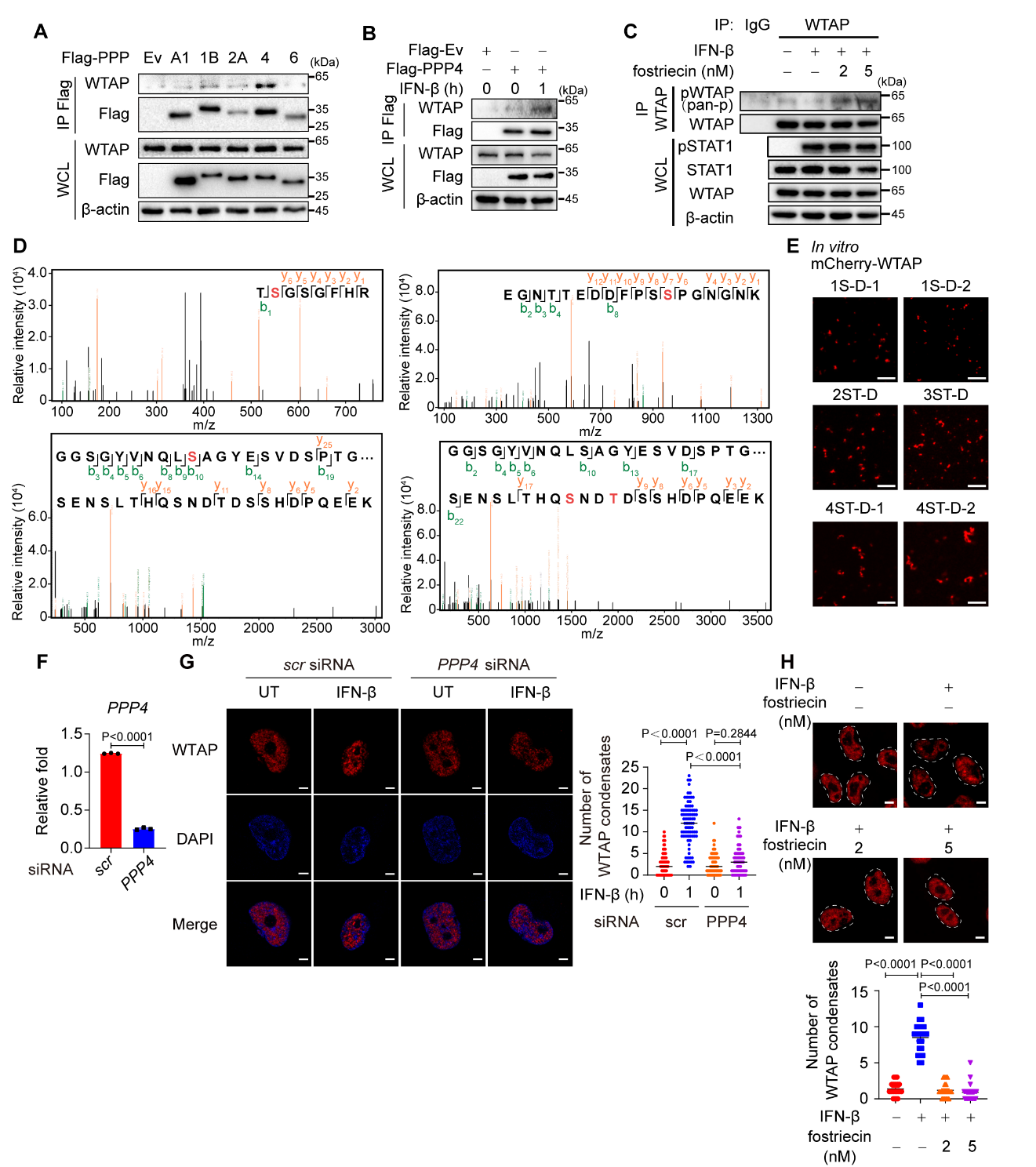

**Figure 2-figure supplement 1. PPP4 is responsible for IFN-β-induced dephosphorylation of WTAP.**

(A) HEK 293T cells transfected with Flag-tagged empty vector (Ev) or protein phosphatases (PPPs). Whole cell lysate (WCL) was collected and immunoprecipitation (IP) experiment with anti-Flag beads was performed, followed with immunoblot.

(B) HEK 293T cells transfected with Flag-tagged Ev or PPP4, followed with IFN-β or left untreated. WCL was collected and IP experiment with anti-Flag antibody was performed, followed with immunoblot.

(C) THP-1-derived macrophages were pre-treated with 2 nM or 5 nM fostriecin for 24 hours or left untreated, followed with 10 ng/mL IFN-β for 1 hour or left untreated. WCL was collected and IP experiment using anti-WTAP antibody or rabbit IgG was performed, followed with immunoblot. pWTAP was detected by anti-phosphoserine/threonine/tyrosine antibody (pan-p).

(D) Data of mass spectrometry (MS) assay of the five phosphorylated sites within WTAP CTD.

(E) Representative images of the phase-separated mCherry-WTAP mutants (containing different numbers of serine (S)/threonine (T) to aspartic acid (D) mutants). Scale bars indicated 5 μm.

(F) THP-1-derived macrophages were transfected with *scramble* (*scr*) or *PPP4*-targeted siRNA. Expression of PPP4C was detected by quantitative real time-polymerase chain reaction (qPCR) assay after siRNA transfection for 48 hours.

(G) THP-1-derived macrophages were transfected with *scramble* (*scr*) or *PPP4*-targeted siRNA and stimulated with or without 10 ng/mL IFN-β for 1 hour. Endogenous WTAP was stained and imaged using confocal microscopy. The number of WTAP condensates that diameter over 0.4 μm of n = 80 cells were counted through ImageJ and shown. Scale bars indicated 5 μm.

(H) mCherry-WTAP-rescued HeLa cells were pre-treated with 2 nM or 5 nM fostriecin for 24 hours or left untreated, followed with 10 ng/mL IFN-β for 1 hour or left untreated. Phase separation of mCherry-WTAP was observed through confocal microscopy. The number of WTAP condensates that diameter over 0.4 μm of n = 20 cells were counted through ImageJ and shown. Scale bars indicated 5 μm.

All error bars, mean values ± SD, P-values were determined by two-way ANOVA test of n = 80 cells in (G) and unpaired two-tailed Student’s *t*-test of n = 20 cells in (H). All error bars, mean values ± SEM, P-values were determined by unpaired two-tailed Student’s *t*-test of n = 3 independent biological experiments in (F). For (A-C, E and G-H), similar results were obtained for three independent biological experiments.

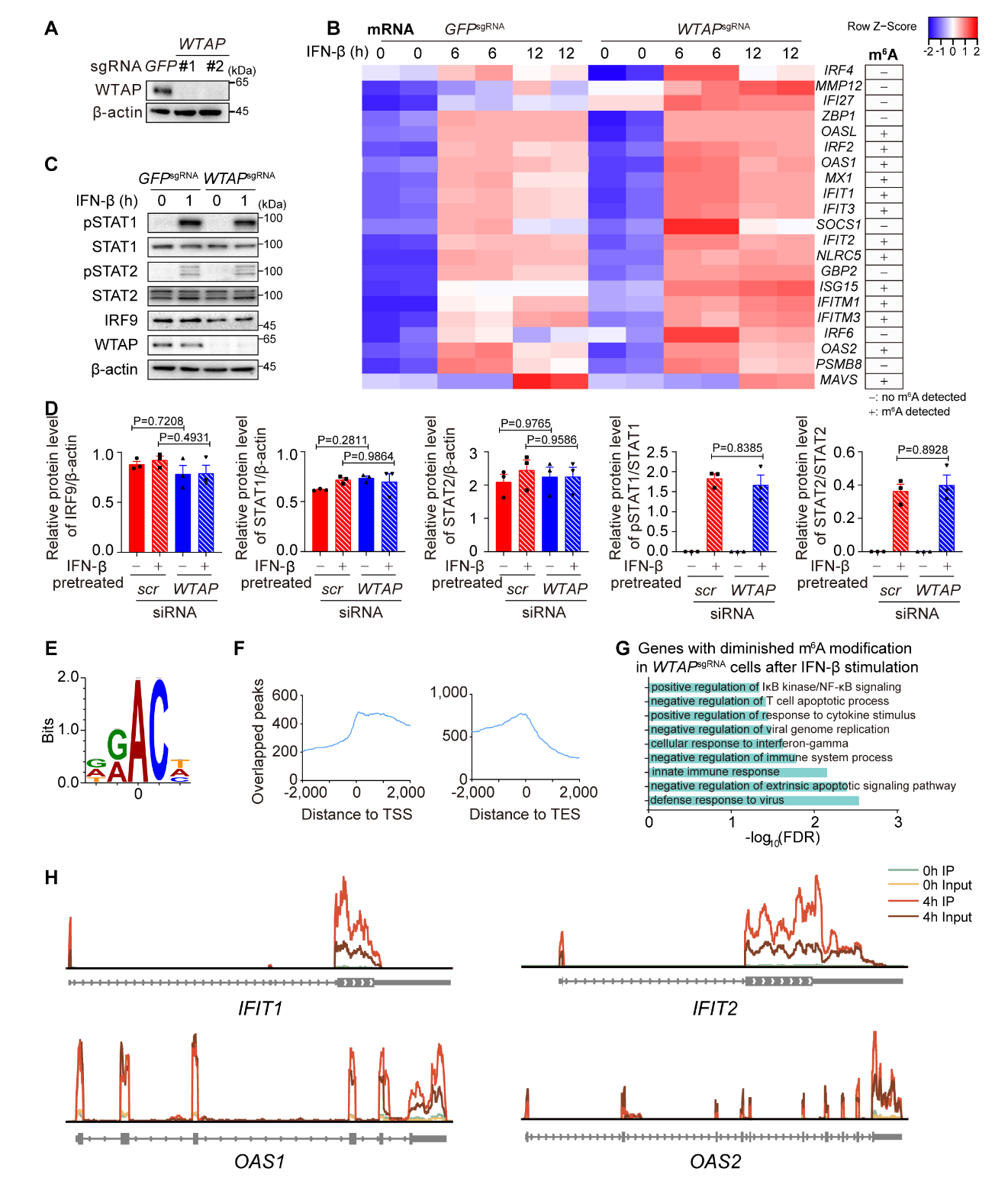

**Figure 3-figure supplement 1. WTAP controls the m^6^A modification of ISG mRNAs.**

(A) *WTAP* knock-down efficiency of control (*GFP*^sgRNA^) and *WTAP*^sgRNA^ #1 or #2 THP-1 was tested by immunoblot.

(B) Transcriptome sequencing analysis of control (*GFP*^sgRNA^) and *WTAP*^sgRNA^ #2 THP-1-derived macrophages stimulated with 10 ng/mL IFN-β for 0, 6, 12 hours, followed with RNA-seq. The count-per-million (CPM) value of core ISGs that upregulated in *WTAP*^sgRNA^ THP-1 cells was drawn by Heatmapper and clustered using Centroid Linkage approach. Control (*GFP*^sgRNA^) and *WTAP*^sgRNA^ #2 THP-1-derived macrophages were treated with 10 ng/mL IFN-β for 4 hours, and m^6^A-modified ISGs analyzed by MeRIP-seq were shown. -: no m^6^A modification detected. +: m^6^A modification detected.

(C-D) Control (*GFP*^sgRNA^) and *WTAP*^sgRNA^ #2 THP-1-derived macrophages were treated with 10 ng/mL IFN-β for 1 hour or left untreated. Whole cell lysate (WCL) was collected followed with immunoblot in (C), while relative protein level and phosphorylation level were measured and analyzed in (D).

(E) Motif enrichment of m^6^A sites were analyzed in mRNAs of the IFN-β-induced WTAP-dependent m^6^A-modified genes through MeRIP-Seq. The position of the methylated adenosine is indicated as zero point.

(F) Distance of m^6^A sites from translation start sites (TSS), translation end sites (TES) within the IFN-β-induced WTAP-dependent m^6^A modified genes were calculated and shown (zero point indicated position of the TSS or TES sites).

(G) Gene ontology analysis for the WTAP down-regulated ISGs with lower m^6^A level in *WTAP*^sgRNA^ #2 cells.

(H) Distribution of m^6^A modification of *IFIT1*, *IFIT2*, *OAS1* and *OAS2*.

All error bars, mean values ± SD, P-values were determined by unpaired two-tailed Student’s *t*-test of n = 3 independent biological experiments in (D). For (A and C), similar results were obtained for three independent biological experiments.

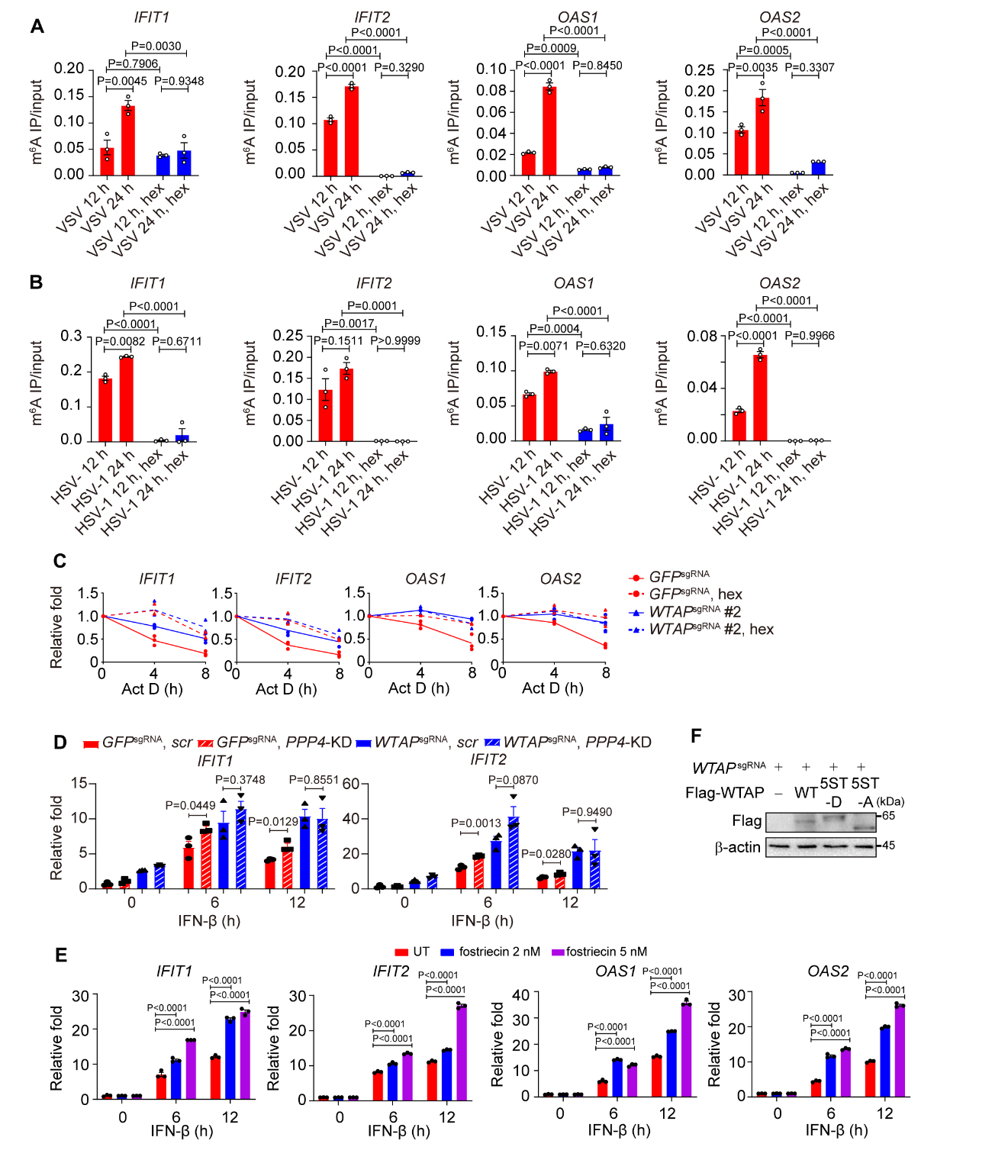

**Figure 4-figure supplement 1.** **Liquid-phase separation of WTAP is crucial for destabilization of ISG mRNAs.**

(A-B) THP-1-derived macrophages were infected with Vesicular Stomatitis Virus (VSV) (m.o.i. = 0.1) (A) or Herpes Simplex Virus-1 (HSV-1) (m.o.i. = 1) (B) for indicated time points, along with or without 5% 1,6-hexanediol (hex) and 20 μg/mL digitonin for 2 hours. MeRIP-qPCR assay of *IFIT1*, *IFIT2*, *OAS1* and *OAS2* was performed and ratios between m^6^A-modified mRNA and input were shown (m^6^A IP/input).

(C) Control (*GFP*^sgRNA^) and *WTAP*^sgRNA^ #2 THP-1-derived macrophages were treated with 10 ng/mL IFN-β together with or without 5% hex and 20 μg/mL digitonin for 2 h or left untreated. After IFN-β treatment, medium with stimuli was replaced by 5 μM actinomycin D (Act D) for indicated time points. RNA was collected and detected by qPCR assay.

(D) Control (*GFP*^sgRNA^) and *WTAP*^sgRNA^ THP-1-derived macrophages were transfected with *scramble* (*scr*) or *PPP4*-targeted siRNA. Cells were treated with 10 ng/mL IFN-β for indicated time points. Expressions of *IFIT1* and *IFIT2* mRNA were detected by qPCR assay.

(E) THP-1-derived macrophages were pre-treated with 2 nM or 5 nM fostriecin for 24 hours or left untreated, followed with 10 ng/mL IFN-β for indicated time points. Expression of *IFIT1*, *IFIT2*, *OAS1* and *OAS2* mRNA was analyzed through qPCR assays.

(F) Immunoblot analysis of the expression of wild type (WT) WTAP,5ST-D or 5ST-A mutant in WT WTAP, 5ST-D or 5ST-A mutant-rescued *WTAP*^sgRNA^ THP-1-derived macrophages .

All error bars, mean values ± SEM, P-values were determined by two-way ANOVA test of n = 3 independent biological experiments in (A-E). For (F), similar results were obtained for three independent biological experiments.

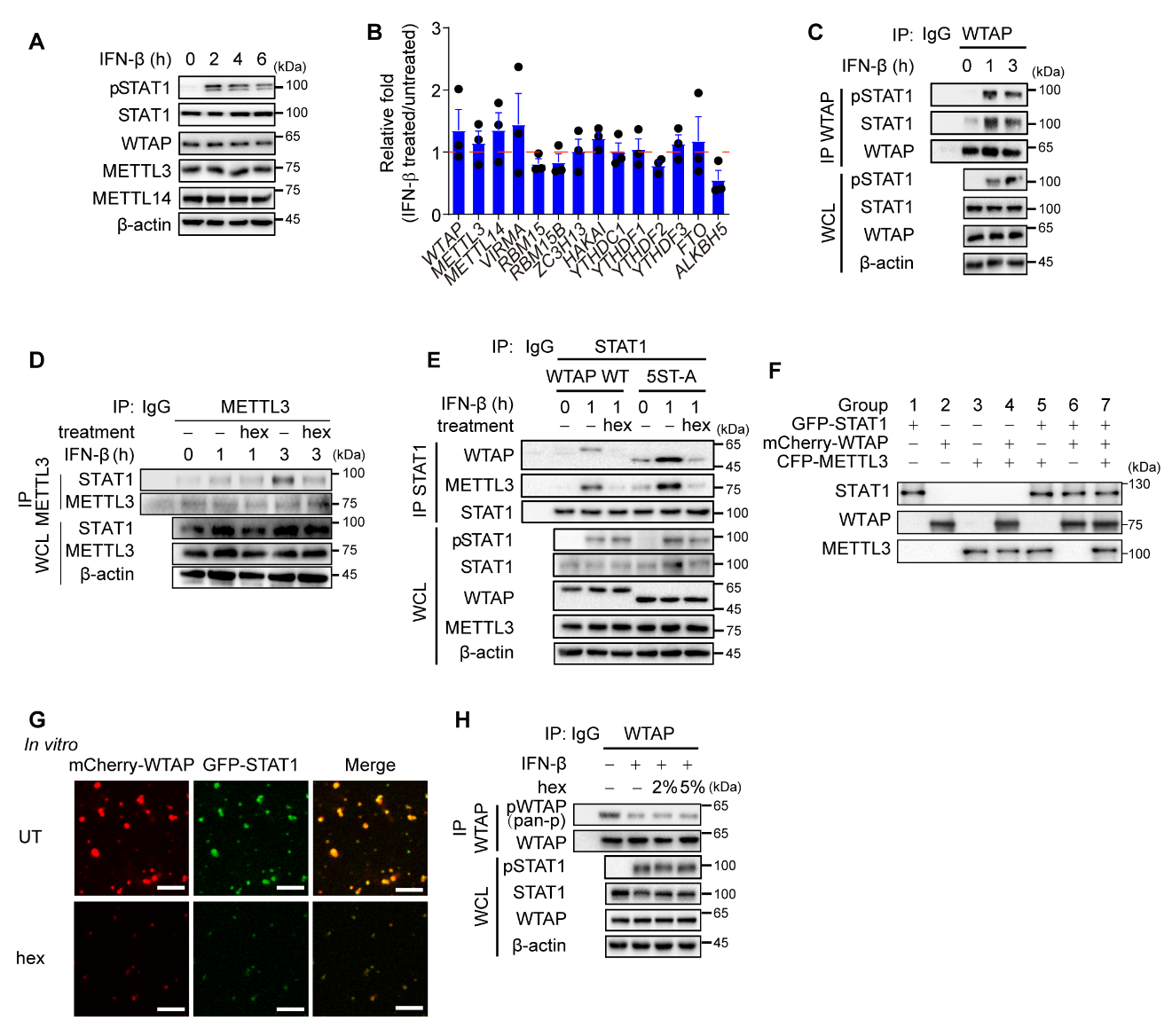

**Figure 5-figure supplement 1. Liquid-phase separated WTAP promotes the interaction between STAT1 and METTL3.**

(A) THP-1-derived macrophages were treated with 10 ng/mL IFN-β for indicated time points. Expression of indicated proteins were detected through immunoblot.

(B) THP-1-derived macrophages were treated with 10 ng/mL IFN-β for indicated time points. Expressions of components of m^6^A methyltransferase complex, m^6^A readers and m^6^A demethylase mRNA were detected through quantitative real-time polymerase chain reaction (qPCR) assay.

(C) THP-1-derived macrophages were treated with 10 ng/mL IFN-β for indicated time points. Whole cell lysate (WCL) was collected and immunoprecipitation (IP) experiment using anti-WTAP antibody or rabbit IgG was performed, followed with immunoblot.

(D) THP-1-derived macrophages were treated with 10 ng/mL IFN-β for indicated time points together with or without 5% 1,6-hexanediol (hex) and 20 μg/mL digitonin. WCL was collected and IP experiment with anti-METTL3 antibody or rabbit IgG was performed, followed with immunoblot.

(E) Wild type (WT) WTAP or 5ST-A mutant-rescued *WTAP*^sgRNA^ THP-1-derived macrophages are stimulated with 10 ng/mL IFN-β together with or without 5% hex and 20 μg/mL digitonin for 1 hour or left untreated. WCL was collected and IP experiment using anti-STAT1 antibody or rabbit IgG was performed, followed with immunoblot.

(F) Protein level of different recombinant proteins of different groups in Fig. 5H-I was detected through immunoblot assays.

(G) Recombinant GFP-STAT1 (10 μM) and mCherry-WTAP (10 μM) were incubated with 5% hex or left untreated (UT) in 37℃. Interaction between WTAP and STAT1 was imaged using confocal microscope. Scale bars indicated 10 μm.

(H) THP-1-derived macrophages were treated with 10 ng/mL IFN-β together with or without 2% or 5% hex and 20 μg/mL digitonin for 1 hour or left untreated. WCL was collected and IP experiment using anti-WTAP antibody or rabbit IgG was performed, followed with immunoblot. pWTAP was detected by anti-phosphoserine/threonine/tyrosine antibody (pan-p).

All error bars, mean values ± SEM, P-values were determined by unpaired two-tailed Student’s *t*-test of n = 3 independent biological experiments in (B). For (A, C-H), similar results were obtained for three independent biological experiments.

**Table S1. Usage of antibodies and other reagents used in this study**

| Antibodies | Source | Identifier | Dilution |
| --- | --- | --- | --- |
| Anti-WTAP | Bethyl Lab | #14994 | 1:1000 for IB  1:100 for IP |
| Anti-WTAP | Santa Cruz Biotechnology | # AB310 | 1:150 for IF |
| Anti-m^6^A | Synaptic Systems | #202003 | 1:400 for IP |
| Anti-phospho-STAT1 (Tyr701) | Cell Signaling Technology | #9167 | 1:1000 for IB |
| Anti-STAT1 | Cell Signaling Technology | #14994 | 1:1000 for IB  1:150 for IF  1:100 for IP |
| Anti-phospho-STAT2 (Tyr690) | Cell Signaling Technology | #88410 | 1:1000 for IB |
| Anti-STAT2 | Cell Signaling Technology | #72604 | 1:1000 for IB |
| Anti-IRF9 | Cell Signaling Technology | #76684 | 1:1000 for IB |
| Anti-mCherry | Proteintech | #26765-1-AP | 1:3000 for IB |
| Anti-METTL3 | Proteintech | #15073-1-AP | 1:1000 for IB |
|  |  |  | 1:150 for IF |
| Anti-PPP4 | Proteintech | #10262-1-AP | 1:1000 for IB |
| Anti-phosphoserine/threonine/tyrosine (pan-p) | Invitrogen | **#61-8300** | 1:1000 for IB |
| Goat anti-rabbit IgG (Alexa Fluor 488) | Invitrogen | #A-11034 | 1:500 for IF |
| Goat anti-mouse IgG (Alexa Fluor 568) | Invitrogen | #A-11031 | 1:500 for IF |
| Goat anti-mouse | Invitrogen | #A-16072 | 1:4000 for IB |
| Goat anti-rabbit | Invitrogen | #A-16104 | 1:4000 for IB |
| Anti-b-actin | Sigma-Aldrich | #A1978 | 1:5000 for IB |
| Anti-Flag (M2) | Sigma-Aldrich | #A8592 | 1:3000 for IB |
| Anti-rabbit IgG | Beyotime Biotechnology | #A7016 | / |
| Anti-mouse IgG | Beyotime Biotechnology | #A7028 | / |

**Table S2. Sequence of primers used in this study**

| sgRNA sequence for KO cell line construction | | |
| --- | --- | --- |
| Targets | **Sequence (5’-3’)** | |
| *GFP* | CATGCCGAGAGTGATCCCGG | |
| *WTAP* #1 | GCTGTAGTCCTGCTGGTACT | |
| *WTAP* #2 | AAGTTGTGCAATACGTCCCT | |
| Sequence of siRNA | | |
| Targets | | **Sequence (5’-3’)** |
| *PPP4* | | CGGCUACCUAUUUGGCAGUGA |
| *STAT1* | | GAAAGAGCUUGACAGUAAA |
| Primers for qPCR and RIP-qPCR assay | | |
| Primers | | **Sequence (5’-3’)** |
| qp-*RPL13A*-F | | GCCATCGTGGCTAAACAGGTA |
| qp-*RPL13A*-R | | GTTGGTGTTCATCCGCTTGC |
| qp-*IFIT1*-F | | TCAGGTCAAGGATAGTCTGGAG |
| qp-*IFIT1*-R | | AGGTTGTGTATTCCCACACTGTA |
| qp-*IFIT2*-F | | GGAGGGAGAAAACTCCTTGGA |
| qp-*IFIT2*-R | | GGCCAGTAGGTTGCACATTGT |
| qp-*OAS1*-F | | AACTGCTTCCGACAATCAAC |
| qp-*OAS1*-R | | CCTCCTTCTCCCTCCAAAA |
| qp-*OAS2*-F | | ACCCGAACAGTTCCCCCTGGT |
| qp-*OAS2*-R | | ACAAGGGTACCATCGGAGTTGCC |
| qp-*METTL3*-F | | TTGTCTCCAACCTTCCGTAGT |
| qp*-METTL3*-R | | CCAGATCAGAGAGGTGGTGTAG |
| qp-*METTL14*-F | | GAACACAGAGCTTAAATCCCCA |
| qp-*METTL14*-R | | TGTCAGCTAAACCTACATCCCTG |
| qp-*RBM15*-F | | ATGCCTTCCCACCTTGTGAG |
| qp-*RBM15*-R | | TCAACCAGTTTTGCACGGAC |
| qp-*RBM15B*-F | | ACCTGGACCACAGCGTATCT |
| qp-*RBM15B*-R | | GGGTTGCGACCAATCACTC |
| qp-*ZC3H13*-F | | GTGCCGTAACTGGCTGAAGA |
| qp-*ZC3H13*-R | | CCTTTACCACGAGGTGAAGGG |
| qp-*VIRMA*-F | | ATACTGATGGTCTGGTGCTAAGA |
| qp-*VIRMA*-R | | TGGAGGGCTTCCATTAAACTGAT |
| qp-*HAKAI*-F | | TGTTACCCGTGCTTCACTTGA |
| qp-*HAKAI*-R | | CTTTGGCGGAATATGGCTCAT |
| qp-*YTHDF1*-F | | ACCTGTCCAGCTATTACCCG |
| qp-*YTHDF1*-R | | TGGTGAGGTATGGAATCGGAG |
| qp-*YTHDF2*-F | | AGCCCCACTTCCTACCAGATG |
| qp-*YTHDF2*-R | | TGAGAACTGTTATTTCCCCATGC |
| qp-*YTHDF3-*F | | TCAGAGTAACAGCTATCCACCA |
| qp-*YTHDF3-*R | | GGTTGTCAGATATGGCATAGGCT |
| qp-*YTHDC1*-F | | AACTGGTTTCTAAGCCACTGAGC |
| qp-*YTHDC1-*R | | GGAGGCACTACTTGATAGACGA |
| qp-*FTO*-F | | AACACCAGGCTCTTTACGGTC |
| qp-*FTO*-R | | TGTCCGTTGTAGGATGAACCC |
| qp-*ALKBH5*-F | | CGGCGAAGGCTACACTTACG |
| qp-*ALKBH5*-R | | CCACCAGCTTTTGGATCACCA |
| Primers for MeRIP-qPCR assay | | |
| Primers | | **Sequence (5’-3’)** |
| MeRIP-*IFIT1*-F | | GGCAGAAGCCCAGACTTACC |
| MeRIP-*IFIT1*-R | | GGGTCCACTTCAAGCACCTT |
| MeRIP-*IFIT2*-F | | GAGATGTGGTGCCCACTAGG |
| MeRIP-*IFIT2*-R | | TGGCCAGTGTCTTAGTCCAA |
| MeRIP-*OAS1*-F | | CAACACAGCCCAGGGATTTC |
| MeRIP-*OAS1*-R | | CCAGTAGATGCAGAGTTGCTG |
| MeRIP-*OAS2*-F | | TCACCCTAGCCCCGTACTTT |
| MeRIP-*OAS2*-R | | AATGAATCGTGACCCCAGCA |
| Primers for ChIP-qPCR assay | | |
| Primers | | **Sequence (5’-3’)** |
| ChIP-*IFIT1*-F | | ATGGTTGCAGGTCTGCAGTT |
| ChIP-*IFIT1*-R | | TGTAAGCTGTGGGTGTGTCC |
| ChIP-*IFIT2*-F | | CCGATGAAACATCCCTCTCTGC |
| ChIP-*IFIT2*-R | | CAAGTGGCCTCTGGTTCCTTTTTG |
| ChIP-*OAS1*-F | | GCAGTGGGCATGTATTGCTG |
| ChIP-*OAS1*-R | | CAGCCCAGCCTTTCTCTGAA |
| ChIP-*OAS2*-F | | ACCCGAACAGTTCCCCCTGGT |
| ChIP-*OAS2*-R | | ACAAGGGTACCATCGGAGTTGCC |
